# Life Cycle Perspectives on Human Health Impacts of Ionic Liquids

**DOI:** 10.1101/091454

**Authors:** Amirhossein Mehrkesh, Arunprakash T. Karunanithi

## Abstract

This study aims to develop a correlative approach to predict the non-cancer human health impacts associated with the direct environmental exposure of common ionic liquids. We assessed the human health impact of these ionic liquids through the integration of the USEtox model with toxicity data and fate and transport parameters. For the first time, we report non-cancer human health characterization factors for commonly used ionic liquids. On the one hand, literature related to environmental aspects of ionic liquids either promotes their environmentally friendly green aspects due to their negligible volatility (no air emissions). On the other hand, a great deal of literature promotes their non-green aspects due to the high toxicity values of certain ionic liquids towards living organisms. In this study, we attempt to integrate these two different diverging opinions to look at the concept of the greenness of ionic liquids from a larger point of view (i.e. from a life cycle assessment perspective).

## 1. Introduction

In recent years, the focus on finding new applications for Ionic liquids(ILs) has increased significantly. Ionic Liquids are, a relatively new class of chemicals, show unique properties such as negligible volatility, non-flammability, high thermal stability, and wide electrochemical windows. These unique and promising characteristics make ionic liquids good candidates for a variety of applications including chemical synthesis^1^, solvent extraction,^2,3^ electrochemistry^4^,carbon capture^5^, cellulose processing^6^, thermal energy storage^7,8^, biomechanics^9^, microfluidics^10^, and many more.

Published literature on ionic liquids has grown exponentially since 1999. New applications are being found for ionic liquids, which has helped these newer compounds find theirway into the industrial applications.^11^ Ionic Liquids are often considered as being inherently green solvents, which have the potential to replace the traditional organic solvents in several applications, mostly due to their non-volatile nature. This advantageous property limits their direct impact on air quality by reducing their emissions to the atmosphere. However, during the past few years, several research studies have countered the view of ionic liquids being all green, and instead reported that many ionic liquids unfortunately shown high toxicity values toward living organisms such as mammals and human cell lines (e.g. HeLa cells).^12^ Even though we believe that both contradictory arguments have their own merits, a realistic picture can certainly be drawn up through the analysis of the human health impacts associated with the life cycle of ionic liquids (from the production of precursor materials to the point of release of ILs into the environment).

Ionic liquids are not yet produced on a large commercial scale, but since the field is developing very rapidly, it is essential to consider their environmental impacts (such as human health impacts) at the design stage rather than after their production.

Research has been focused on toxicity of ionic liquids and their impacts on human health and ecology.^13–15^ Several research have studied the toxicity of ionic liquids towards different living organisms including bacteria such as E.coli,^16,17^ and Vibrio fischeri^18^, microalga such Pseudokirchneriella subcapitata (PKS), planktons such as Daphnia magna and fish such as Danio rerio (zabra fish).^19^ However, there is still a long way to have a universal knowledge on the potential hazard, bioaccumulation, biodegradability and eco-toxicity of ionic liquids as well as their impact on the human health.^13–15,20–21^ To date, the effect of room temperature ionic liquids (RTILs) on some microorganisms and human cell lines has been studied. Ganske et al. showed in their paper that RTILs can potentially have significant inhibitory effects on the growth of microorganisms such as E.coli, Pichiapastoris and Bacilluscereus.16 Among the published literatureon the topicof toxicity of ionic liquids(ILs), only fewa have studied the toxicity of RTILs on human cell lines, such as prototypical human epithelial Caco-2, HeLa cells (an immortal human cell line), and human colon carcinoma HT-29 cell lines.^22,23^ Since potentially a wide variety of ionic liquids consisting of different cations, anion and side chain groups can be synthetized, it is important to develop correlative or QSAR models to predict the toxicity of ionic liquids towards different organisms. This study aims to develop a correlative model to predict the non-cancer human health toxicity of ionic liquids, ED50, from their in-vitro cytotoxicity data.

## 2. Methods

Two correlative approaches were used to convert the in-vitro (toxicity towards microorganisms) cytotoxicity data(IC_50_) to in-vivo toxicity(toxicity towards mammalian species such as rat and mouse) data (LD_50_) and in the next step another correlation was used to predict in-vivo toxicity data to non-cancer human heath toxicity.

An EPA document reported a correlation model (Multicentre Evaluation of In-Vitro Cytotoxicity) along with a methodology developed foraselect number of chemical compounds to predict the in-vivo toxicity values (LD_50_) from in-vitro cytotoxicity data.^24^ In this study, we improved this general correlation by using more categorized chemicals resulting in having abetterthe correlation coefficient,R2, as shown in Fig.1.

**Fig.1.**
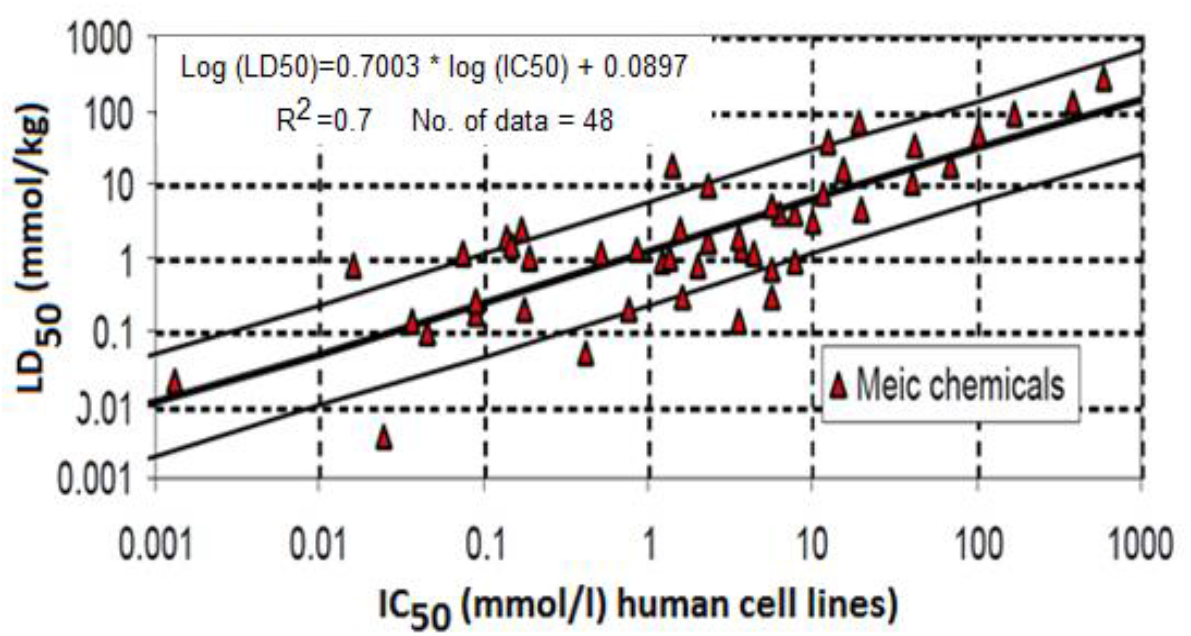
A correlative linearmodel of LD_50_ toxicity data

LD_50_ or the lethal dose of 50%, is the amount of a chemical which kills 50% of the population of a test species (used in toxicity evaluations) exposed to that. The EPA study used IC_50_ toxicity data of selected chemicals towards HeLa cell lines to predict their in-vivo (LD_50_) toxicity values. IC_50_ is an index of the toxicity of a substance defined as the concentration of that chemical which have inhibitory effects on the growth of the 50% of the test population. HeLa cell used in this study is an immortal cell line,which is the oldest yet most commonly used human cell line forscientificresearch.^25^ In the next step, we explain how LD_50_ data predicted from the EPA's correlative model can be further used to estimate the human health impacts of ionic liquids.

### 2.1. Life Cycle Assessment (LCA)

As it was mentioned earlier, to have a realistic perception on the greenness of a chemical compound, a life cycle assessment is needed as a comprehensive and reliable approach towards the quantification of the compound's greenness.

LCA is a method ortechnique used to assess the environmental impacts associated with the lifecycle of a product, process, or service through, compiling a general inventory including relevant energy/material inputs and environmental emissions (releases), evaluating/weighing the environmental impacts potentially associated with identified inputs (materials and energy) and outputs (emissions), and interpreting the results achieved in previous steps to make informed decisions.

A life cycle assessment (LCA) study typically consists of four distinguished steps as the following:

1) Goal and scope definition, 2) Inventory analysis, 3) Impact assessment, and 4) Interpretation as shown in Fig.2.

**Fig.2.**
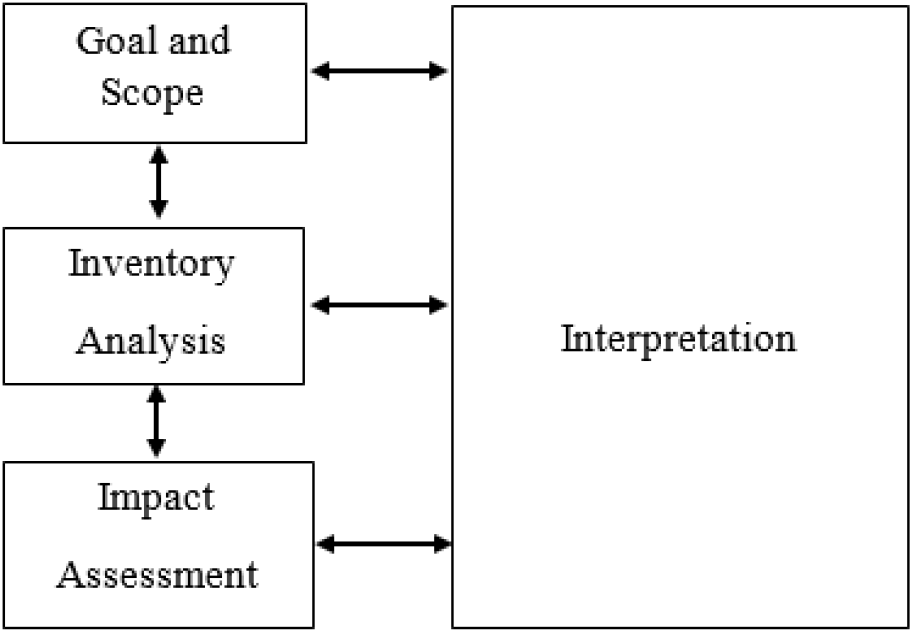
Framework of a life cycle assessment study

#### 2.1.1 Environmental Impact Assessment (EIA)

One of the stages in a lifecycle assessment is an environmental impact assessment (EIA). EIA is the method of measuring the anticipated impacts of a proposed process on the environment. In the case of EIA showing unacceptable effects, design features or further mitigation methods can be used to reduce or even avoid the impacts.

The U.S. environmental protection agency, has introduced TRACI, the Tool for the Reduction and Assessment of Chemical and other environmental Impacts, to assist in life cycle assessment, industrial ecology, pollution prevention and process design.

To develop TRACI, the impact assessment methodology needed to be consistent with previous modeling especially of EPA. The human health non-cancer and cancer categories were mostly based on the assumptions made in EPA's Exposure Factors Handbook and Risk Assessment Guidance for Superfund. TRACI has listed many characterization factors for different category of environmental impacts and different route of exposures (ecotoxicity, human health, global warming etc.) for more than 5000 chemicals. Environmental impact assessments can be done using characterization factors listed in TRACI. This can be done by a weighted as follows:

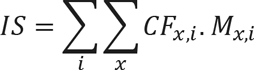

where “IS” being the impact score (e.g. human health toxicity (cases); CF_x,i_ is the characterization of chemical x which is released to compartment i (cases/kg) and M_x,i_ is the emission of substance x to compartment i (kg). The two summations ensure that all the impacts associated with the release of different substances into different compartments were considered to calculate the total impact factor. The USEtox™ model^26^ calculates awide variety of characterization factors for both carcinogenicand non-carcinogenicimpacts of chemical emissions to different environmental compartments (urban air,rural air, freshwater, sea water …). Table 1 tabulated the characterization factors related to the human health impacts of selected chemicals. The characterization factor units for the human toxicity is cases/kg_emission_. which is summarized as CTU (Comparative Toxic Unit) to emphasize on the comparative nature of CFs.

**Table 1.** Characterization factors associated with human health impacts (TRACI database)

| Substance Name | Human health CF<br>[CTU<sub>noncancer</sub>/kg],<br>Emission to urban<br>air, non-canc. | Human health CF<br>[CTU<sub>noncancer</sub>/kg],<br>Emission to cont.<br>freshwater, non-canc. | Human health CF<br>[CTU<sub>noncancer</sub>/kg],<br>Emission to cont.<br>natural soil, non-canc. | CF Flag HH<br>non-carcinogenic |
| --- | --- | --- | --- | --- |
| FORMALDEHYDE | 2.67E-07 | 3.24E-07 | 9.96E-08 | Recommended |
| PHENOBARBITAL | n/a | n/a | n/a | n/a |
| CYCLOPHOSPHAMIDE | n/a | n/a | n/a | n/a |
| PREDNISOLONE | n/a | n/a | n/a | n/a |
| ESTRADIOL | n/a | n/a | n/a | n/a |
| P,P'-DDT | 4.65E-05 | 4.25E-04 | 9.67E-07 | Recommended |

The effective dose for non-cancer human health (ED_50_) predicted through the correlation developed in this study can be used to complete the necessary data for doing a comprehensive life cycle assessment (LCA) on different ionic liquids.

#### 2.1.2. Characterization factor (CF) development

Life cycle impact assessment (LCIA) methods use characterization factors to quantify the extent to which a chemical substance contributes to the environmental impacts. In this work the USEtox model^26^, which is a state-of-the-art modeling framework based on scientific consensus for characterizing human and ecotoxicological impacts of chemicals, was utilized to develop characterization factors. In USEtox, the human health characterization factors of a substance of interest can be estimated through incorporating three parameters, as shown in Eqn. 1.

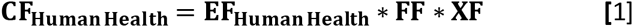

where EF relates to the inherent toxicity, FF relates to the fate factor and XF relates to potential routes and amount of exposure to the chemical of interest, respectively. In the following sections, the procedure for the development of each of these factors will be explained.

##### Fate Factor

Multimedia fate and transport models as shown in Fig.3 are utilized to derive the environmental fate factors. These models represent the environment as several homogeneous compartments(e.g. air, water, and soil). The Intermedia transfer and distribution of a chemical compounds between different environmental compartments are modeled as a set of mass balance equations assuming equilibrium conditions. The fatefactor, representing thepersistence of asubstancein an environmental compartment (residence time in days) depends on biodegradability, physicochemical properties and partitioning coefficients of the substance of interest. The current study estimates the fate factor of different ionic liquids through the approach proposed by USEtox model.

**Fig.3.**
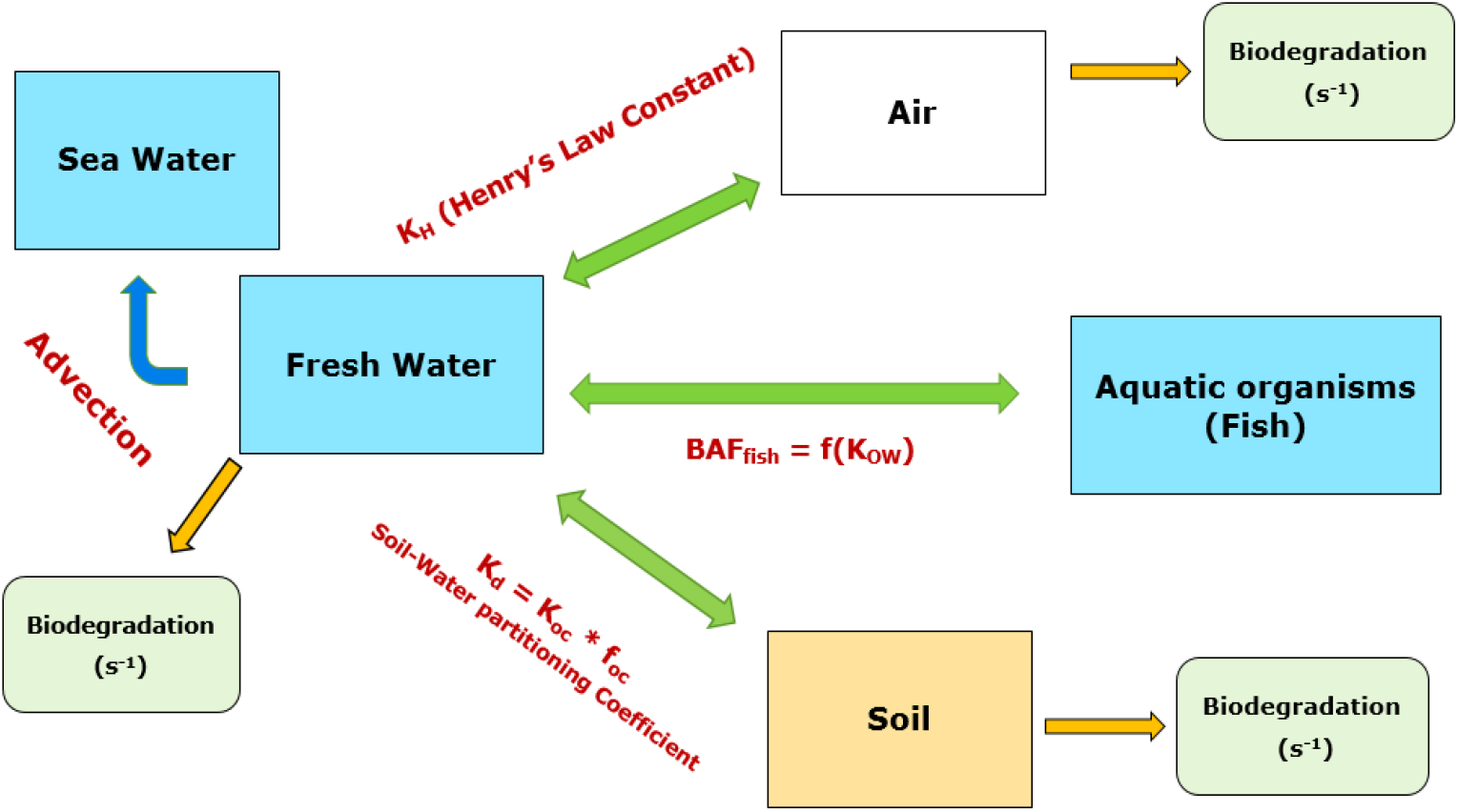
Multimedia Fate and transport model

##### Exposure Factor

In USEtox, the exposure factors related to human health reflect the rate at which a pollutant can transfer from a receiving compartment into the human population through a series of exposure pathways. In USEtox™, air (inhalation), drinking water (ingestion), unexposed produce (below-ground root crops), exposed produce (above-ground leaf crops, e.g. fruit and cereals), meat, dairy products and Fish are the pathways considered for the human exposure to chemical pollutants.

The exposure factor(days-1) fordrinking water ora specificfood item at a certain scale equals.

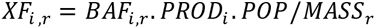

where BAF_i,r_ being the bioaccumulation factor of the chemical of interest of exposure pathway i (e.g. fish) through compartment r (e.g. seawater or freshwater) in kg/kg, PRODi is the production of item *i* in the exposure pathwayperperson (suggestedvalueof 0.04-0.045 kg/day/person for fish living in freshwater), the variable POP represents the population number(e.g. 900 million on the continental scale), and MASSr being the mass of compartment r (e.g. 6.8×10^14^ kg of continental freshwater).

An illustrative example of the calculation of exposure factor(XF) for a specific ionic liquid, [Bmim]^°^[PF_6_]-. The ionic liquid, Bmim PF6 has a BAF of 5.73×10^-3^ kg/kg, PROD_i_ is assumed to be equal to 0.04 kg/day/person for freshwater fish, POP is 900 million on the continental and MASSr=6.8×10^14^ kg for continental freshwater.

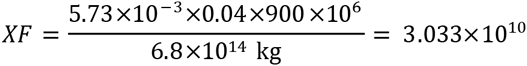

This approach was used to calculate the exposure factors for some common ionic liquids as listed in Table 2.

**Table 2.** Exposure factors of some common ionic liquids

| IL | BAF<sub>i,r</sub> (kg/kg)<sup>30</sup> | XF<sub>i,r</sub> |
| --- | --- | --- |
| [Bmim]<sup>+</sup>[Br]<sup>-</sup> | 1.58×10<sup>-4</sup> | 8.15×10<sup>-12</sup> |
| [Bmim]<sup>+</sup>[Cl]<sup>-</sup> | 8.55×10<sup>-4</sup> | 4.53×10<sup>-11</sup> |
| [Bmim]<sup>+</sup>[BF<sub>4</sub>]<sup>-</sup> | 8.94×10<sup>-4</sup> | 4.73×10<sup>-11</sup> |
| [Bmim]<sup>+</sup>[PF<sub>6</sub>]<sup>-</sup> | 5.73×10<sup>-3</sup> | 3.03×10<sup>-10</sup> |
| [BPy]<sup>+</sup>[Cl]<sup>-</sup> | 6.04×10<sup>-4</sup> | 3.19×10<sup>-11</sup> |

##### Human Health Effect Factor

In USEtox, the human health effect factors (EF) are used to show how change in life time intake of a pollutant (cases/kg intake) can change the life time disease probability. In USEtox™, distinct effect factors are derived for carcinogenic and non-carcinogenic effects. For each effect type, two different exposure routes of ingestion and inhalation are studied separately. The effect factors (EF) related to the human health are calculated if a linear concentration–response up to the point of 50% life time disease probability, as shown in Fig.4, is in effect.

**Fig.4.**
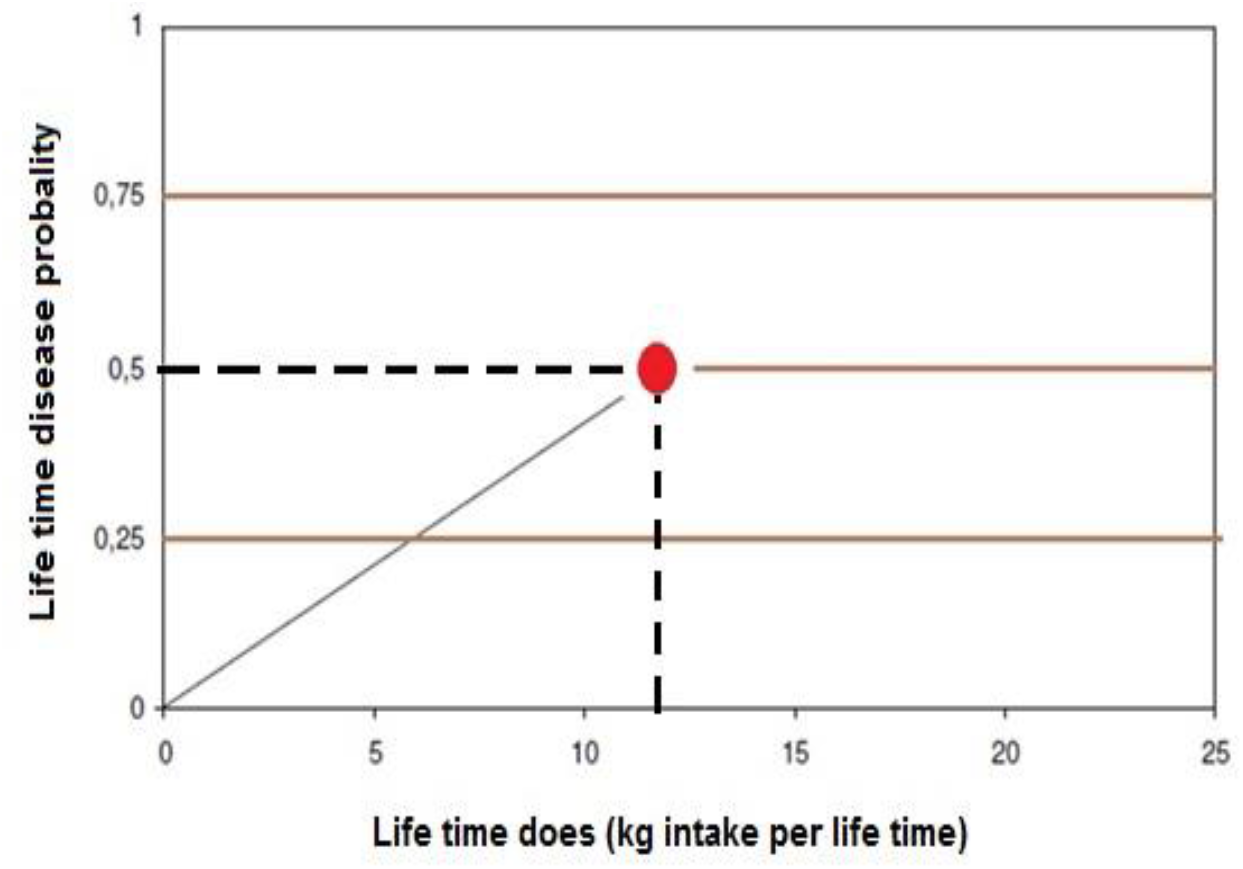
Concentration–response up to the point of 50% life time disease probability

The human health toxicological effect factor of a chemical of interest can be estimated through the following equation:

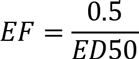

For non-carcinogenic and carcinogenic effects, the ED50_h,j_ for human health related to ingestion or inhalation exposure (kg/person/lifetime) is calculated using the equation below.

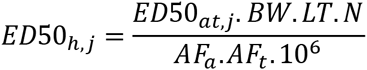

where ED50_a,t,j_ is the daily dose for the test animal (e.g. a rat or a mouse) during the time t (e.g. chronic or sub-chronic) per kg body weight (BW) of the species that causes a 50% chance of acquiring a disease through exposure route of j (mg.kg^-1^.day^-1^). AF_a_ is the extrapolation factor used to convert the results related to one species to the other one (interspecies factors) which are listed in Table 3. AFt is the extrapolation factor to address the differences in time of exposure, (i.e. a factor of 2 is used to convert sub-chronicto chronicexposure)^27^, BWis representing the average body weight of ahuman being (70-75 kg), LT being the average lifetime of humans (70 years) and N is the number of days in one year used for unit conversion (365 days. year^-1^).^26^

**Table 3.** A list of AF_a_, interspecies factors

| Type | $AF_{interspecies}$ | Average<br>Bodyweight (kg) |
| --- | --- | --- |
| Human | 1.0 | 70 |
| Pig | 1.1 | 48 |
| Dog | 1.5 | 15 |
| Monkey | 1.9 | 5 |
| Cat | 1.9 | 5 |
| Rabbit | 2.4 | 2 |
| Hen | 2.6 | 1.6 |
| Mink | 2.9 | 1 |
| Guinea pig | 3.1 | 0.750 |
| Rat | 4.1 | 0.25 |
| Hamster | 4.9 | 0.125 |
| Gerbil | 5.5 | 0.075 |
| Mouse | 7.3 | 0.025 |

In the next step, we chose 38 chemicals for which both the non-cancer human health toxicology data (ED_50_) from TRACI and in-vivo toxicity (LD_50_) data for oral exposure of rats from the literature were available. We tried our best to select at least one chemical compound from each category (acids, aldehydes, aromatics, etc.). A linear correlation was developed to predict the non-cancer human health effective dose of chemicals from their in-vivo toxicicty dataas shown in Fig.5.

**Fig.5.**
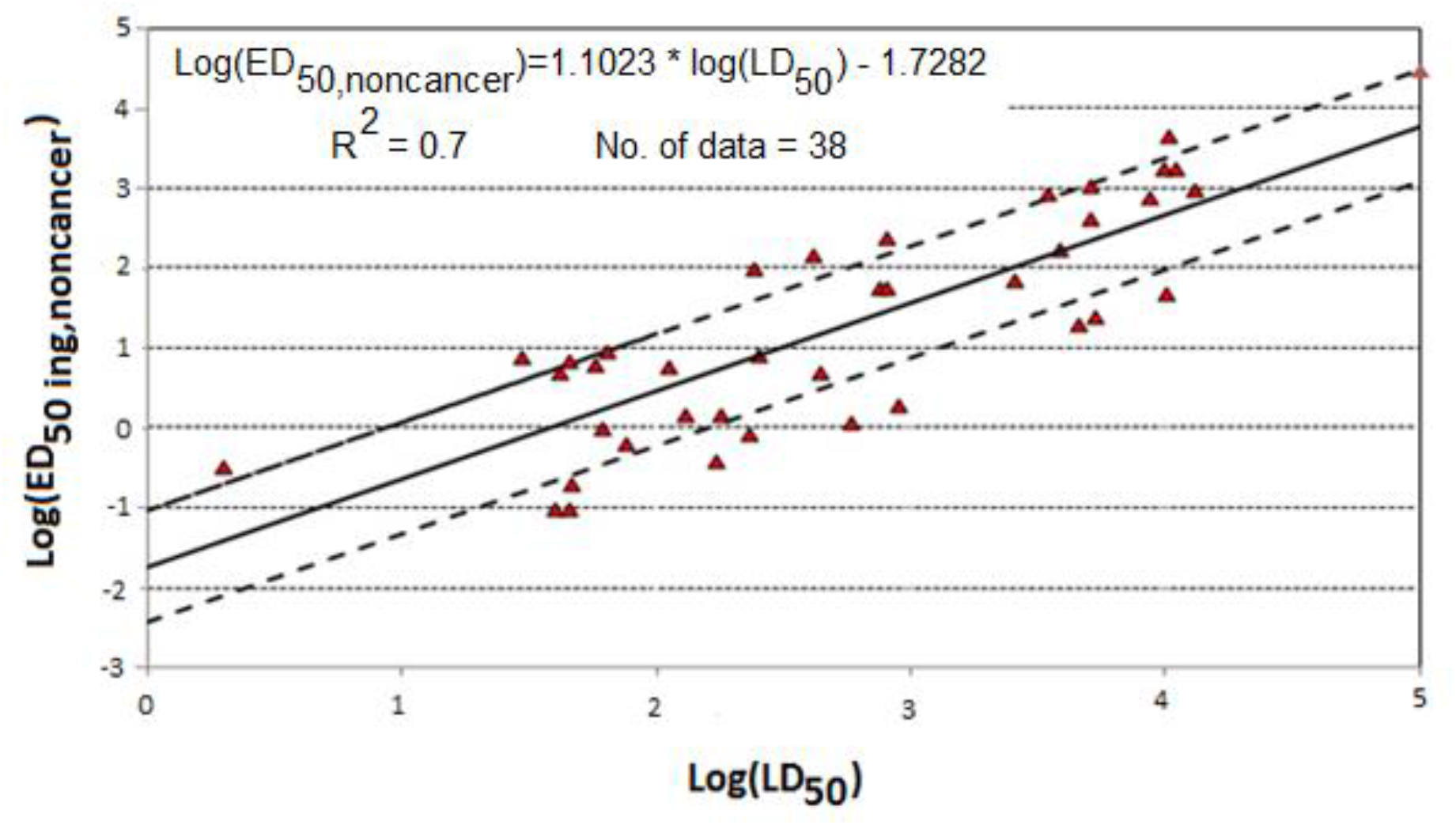
A linearcorrelation for ED_50_ vs. LD_50_ data

The algorithm shown in Fig. 6 can be used for any chemical of interest to give us an approximate value(as an start point) of ED_50,non-cancer_ from the IC_50_ cytotoxicty data of that chemical towards human cell lines (e.g. HeLa cell).

**Fig.6.**
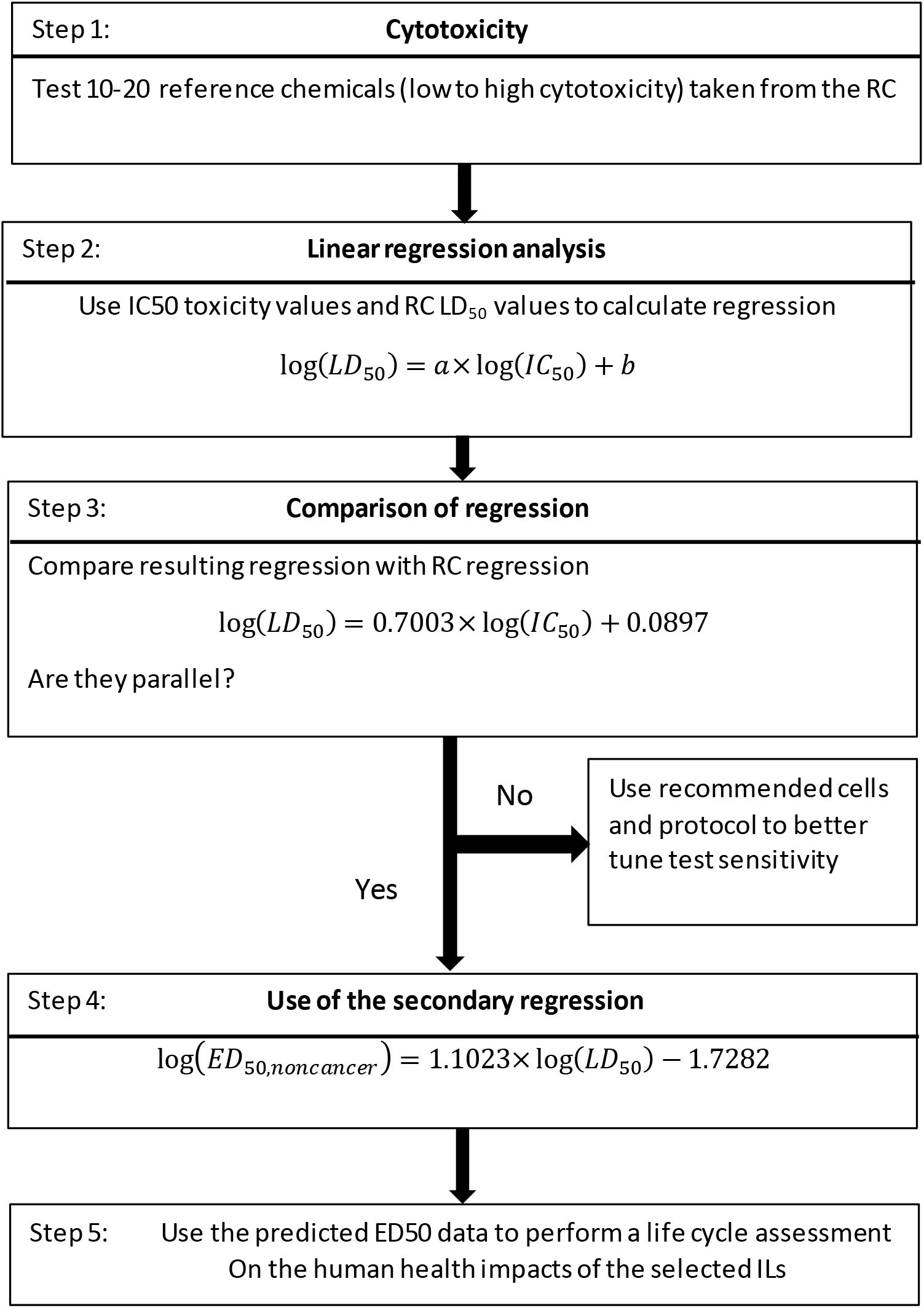
Algorithm to predict the ED50 for a chemical of interesta

The predicted value of in-vivo toxicity (LD_50_) along with the non-cancer human health toxicity (ED_50_) data for a few ionic liquids are listed in Table 4.

**Table 4.** Predicted values of in-vivo toxicity (LD_50_) and non-cancer human toxicity (ED_50_) for ILs

| IL | MW (g/mol) | Log(IC<sub>50</sub>) [μM]<sup>14</sup> | Log(LD<sub>50</sub>) [mmol/kg] | ED<sub>50,ing non-cancer</sub> [kg/lifetime] |
| --- | --- | --- | --- | --- |
| [Bmim]<sup>+</sup> [Br]<sup>-</sup> | 219.122 | 3.44 | 2.498 | 10.6 |
| [Bmim]<sup>+</sup> [Cl]<sup>-</sup> | 174.67 | ----- | ----- | ----- |
| [Bmim]<sup>+</sup> [BF<sub>4</sub>]<sup>-</sup> | 226.03 | 3.72 | 2.694 | 17.43 |
| [Bmim]<sup>+</sup> [PF<sub>6</sub>]<sup>-</sup> | 284.18 | 4.14 | 2.988 | 36.76 |
| [BPy]<sup>+</sup> [Cl]<sup>-</sup> | 171.667 | ----- | ----- | ----- |

Stepnowski et al^28^ reported a value of 13.9 mmol. L^−1^ for IC_50_ of [Bmim]^+^ [PF6]^−^ using of this value in the correlative model developed in this study resulted in an LD_50_ of 276.43mg/kg, which is in close vicinity of the experimental value of 300 mg/kg for the LD_50_ of the selected ionic liquid for oral intake by rats.^29^

In our previous study, the fate factors of common ionic liquids were calculated.^30^ These fate factors, along with the effect and exposure factors calculated in this study, were used to develop new characterization factors fornon-cancerhuman health impacts of common ionic liquids as tabulated in Table 5.

**Table 5.**
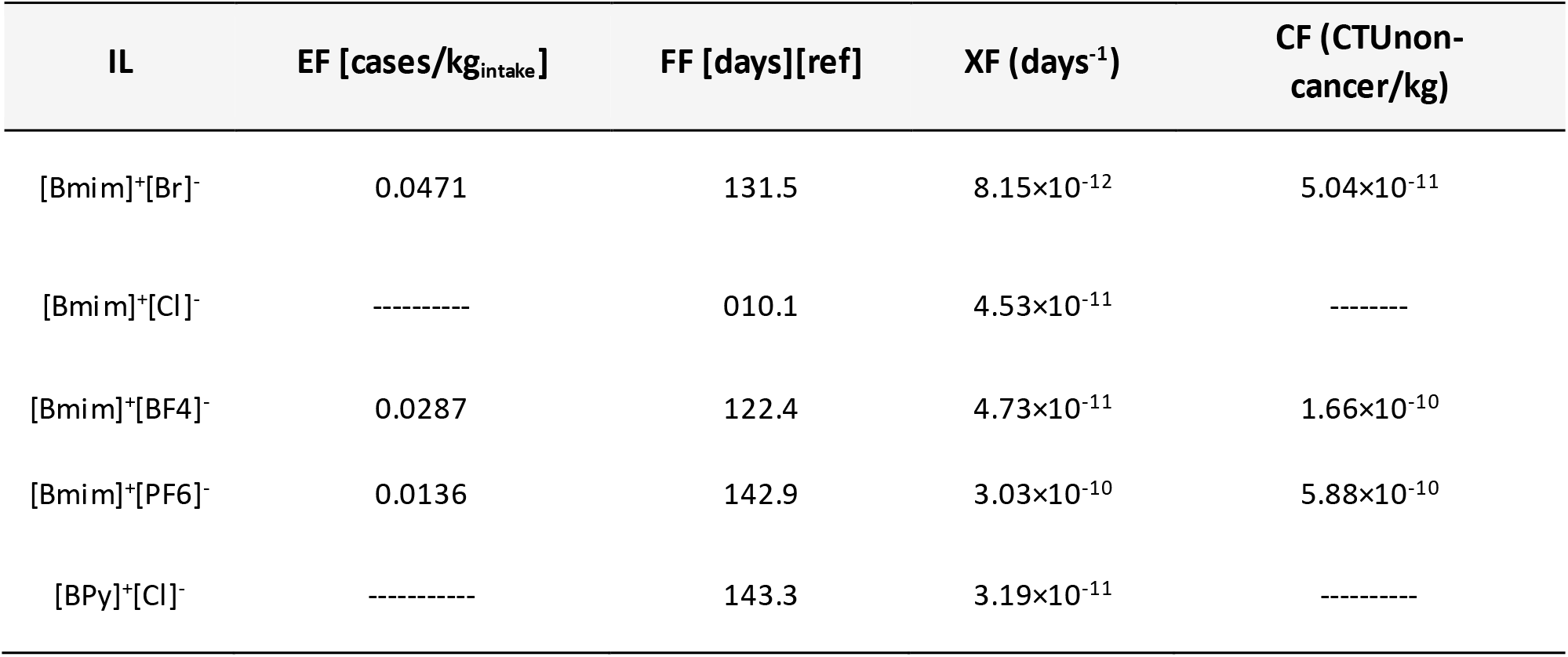
Characterization factors of non-cancer human health of conventional ionic liquids

## Acknowledgement

This material is based upon work supported by the United States National Science Foundation (CAREER Program) under Grant No. 1151182.

